# A deep learning framework for characterization of genotype data

**DOI:** 10.1101/2020.09.30.320994

**Authors:** Kristiina Ausmees, Carl Nettelblad

## Abstract

Dimensionality reduction is a data transformation technique widely used in various fields of genomics research. The application of dimensionality reduction to genotype data is known to capture genetic similarity between individuals, and is used for visualization of genetic variation, identification of population structure as well as ancestry mapping. Among frequently used methods are PCA, which is a linear transform that often misses more fine-scale structures, and neighbor-graph based methods which focus on local relationships rather than large-scale patterns.

Deep learning models are a type of nonlinear machine learning method in which the features used in data transformation are decided by the model in a data-driven manner, rather than by the researcher, and have been shown to present a promising alternative to traditional statistical methods for various applications in omics research. In this paper, we propose a deep learning model based on a convolutional autoencoder architecture for dimensionality reduction of genotype data.

Using a highly diverse cohort of human samples, we demonstrate that the model can identify population clusters and provide richer visual information in comparison to PCA, while preserving global geometry to a higher extent than t-SNE and UMAP. We also discuss the use of the methodology for more general characterization of genotype data, showing that models of a similar architecture can be used as a genetic clustering method, comparing results to the ADMIXTURE software frequently used in population genetic studies.

## INTRODUCTION

The increasing availability of large amounts of data has led to a rise in the use of machine learning (ML) methods in several fields of omics research. For many applications dealing with complex and heterogeneous information, the data-driven approach has become a promising alternative or complement to more traditional model-based methods (Xu and Jackson 2019; Libbrecht and Noble 2015; Schrider and Kern 2018).

Deep learning (DL) is an active subdiscipline of ML that has had a large impact in several fields, including image analysis and speech recognition (LeCun *et al*. 2015). DL methods comprise models that compute a nonlinear function of their input data using a layered structure that learns abstract feature representations in a hierarchical manner, and can be used for supervised learning tasks such as prediction and also in unsupervised settings for pattern recognition and data characterization problems (Goodfellow *et al*. 2016). A key aspect of DL is that the features used in data transformation are learned by the model as opposed to being defined by the researcher, resulting in a higher level of flexibility than alternative ML algorithms such as support vector machines (Zou *et al*. 2019).

Advances have been made in developing DL techniques for various types of omics data (Eraslan *et al*. 2019a). The current state-of-the-art for predicting effects of genetic variants on splicing is a DL model (Cheng *et al*. 2019). The DeepBind model (Alipanahi *et al*. 2015) outperformed several previous non-DL approaches for predicting sequence specificities of DNA-binding proteins. For the task of variant calling of single-nucleotide polymorphisms (SNPs) and small indels, the DeepVariant model of Poplin *et al*. (2018) was shown to give improved results over existing tools (Nawy 2018).

DL has also been applied to unsupervised problems, including imputation of metabolite and SNP data (Scholz *et al*. 2005; Chen and Shi 2019; Sun and Kardia 2008), de-noising of ChIP-sequencing data (Koh *et al*. 2017) and outlier detection of RNA sequencing gene expression data (Brechtmann *et al*. 2018). In the field of single-cell RNA-sequencing, DL methods have been used for imputation, de-noising as well as dimensionality reduction (Talwar *et al*. 2018; Ding *et al*. 2018; Eraslan *et al*. 2019b).

Dimensionality reduction is a data transformation technique that is commonly applied to SNP data in the fields of population and quantitative genetics. Applications include visualization of genetic variation, detection of population structure and correcting for stratification in genome-wide association studies (GWAS) (Patterson *et al*. 2006; Price *et al*. 2006). One of the most widely used methods for performing dimensionality reduction is Principal Component Analysis (PCA), in which a linear transformation is made onto uncorrelated dimensions that maximize the variance of the projected data (Pearson 1901). It has been shown that the lower-dimensional representation resulting from PCA can capture patterns in genetic variation, e.g. by reconstructing geographical relationships from genotype data (Novembre *et al*. 2008).

Although PCA is an efficient and reliable method, there are limitations associated with it. Firstly, it can be sensitive to attributes of sequence data such as the presence of rare alleles and SNPs that are correlated due to linkage disequilibrium (LD), which can cause groupings of samples that reflect such phenomena rather than genome-wide population structure (Ma and Shi 2020; Tian *et al*. 2008). To avoid such spurious effects, a stringent filtering procedure to remove low frequency variation and SNPs in high LD is usually required prior to performing PCA, although methods to handle LD by e.g. shrinkage methods have been proposed (Zou *et al*. 2010). Further limitations of PCA are related to the inability to capture nonlinear patterns in the data, as discussed in e.g. Alanis-Lobato *et al*. (2015), where the nonlinear method of non-centred Minimum Curvilinear Embedding (ncMCE) is proposed and shown to detect population structure in cases where PCA fails.

A family of nonlinear methods that has seen increased use on SNP data is neighbor graph-based models, including t-distributed stochastic neighbor embedding (t-SNE) and Uniform Manifold Approximation and Projection (UMAP) (van der Maaten and Hinton 2008; McInnes *et al*. 2020). These methods consider neighboring samples around each data point and try to find a lower-dimensional representation that preserves the distances between the points in the neighborhood. Both t-SNE and UMAP have been shown to be able to capture more fine-scale population structure than e.g. PCA, but the focus on preserving local topology results in a projection in which distances between larger clusters are more difficult to interpret (Gaspar and Breen 2019; Diaz-Papkovich *et al*. 2021).

More recently, DL methods for dimensionality reduction of SNP data have also been introduced. In Yelmen *et al*. (2021), the focus is on generating artificial genotypes, but one type of model considered, restricted Boltzmann machines (RBMs), projects the data to a reduced-dimensionality space which is compared to that of PCA. A DL approach based on variational autoencoders is presented in Battey *et al*. (2020), where they show that their model can capture subtle features of population structure, while preserving global geometry to a higher degree than both t-SNE and UMAP.

In this paper, we present a DL framework denoted Genotype Convolutional Autoencoder (GCAE) for nonlinear dimensionality reduction of SNP data based on convolutional autoencoders. The main differentiating feature between GCAE and the other DL methods mentioned is that our model makes use of convolutional layers, which take into account the sequential nature of genotype data. We describe adaptations to network architecture implemented to capture local as well as global patterns in sequence data, and compare dimensionality reduction performance to that of PCA, t-SNE and UMAP on a highly diverse cohort of human samples. We also demonstrate the broader applicability of the framework for general characterization of SNP data by showing that minor modifications in network structure can produce a model for solving the genetic clustering problem, and compare results to a model-based method commonly used in population genetic studies.

## MATERIALS AND METHODS

### Model architecture and training strategy

The proposed model for dimensionality reduction of SNP data is a convolutional autoencoder. Autoencoders are a class of DL models that transform data to a lower-dimensional latent representation from which it is subsequently reconstructed (Kramer 1991; Hinton and Salakhutdinov 2006). The idea is to learn features, or variables derived from the original data, that capture the important characteristics of the data. The structure of the model is shown in Figure 2. It comprises four types of layers that transform the input in a sequential manner: convolutional, pooling, fully-connected and upsampling layers.

**Figure 1.**
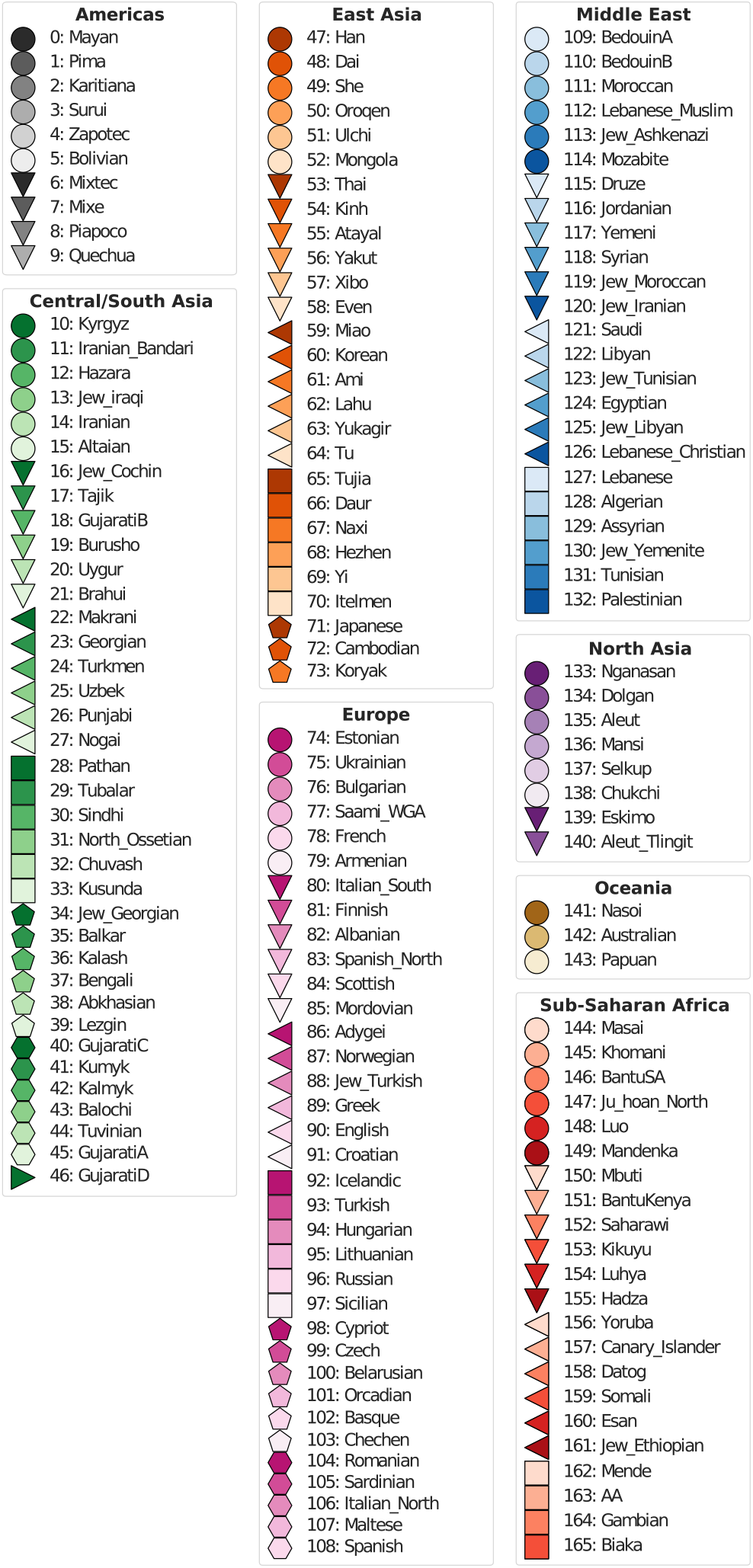
Populations and superpopulations of the Human Origins panel of genotype data. The coloring serves as a legend for Figure 3 and the numbering for Figures 5 and 6.

**Figure 2.**
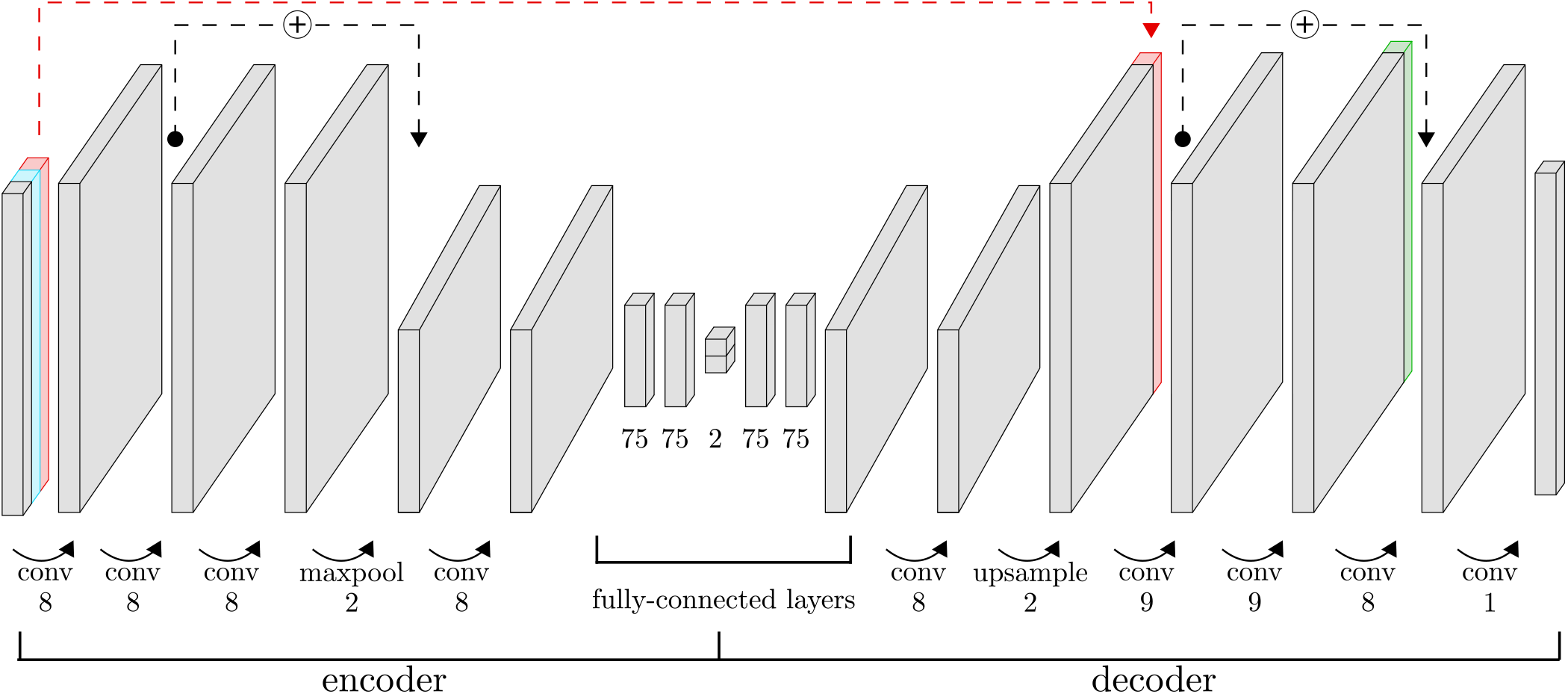
Architecture of the GCAE model used for dimensionality reduction. The encoder transforms data to a lower-dimensional latent representation through a series of convolutional, pooling and fully-connected layers. The decoder reconstructs the input genotypes. The input consists of three layers: genotype data (gray), a binary mask representing missing data (blue), and a marker-specific trainable variable per SNP (red). The red dashed line indicates where this marker-specific variable is concatenated to a layer in the decoder. Another marker-specific trainable variable, shown in green, is also concatenated to the second-last layer in the decoder. Black dashed lines indicate residual connections, where the output of a layer is added to that of another layer later in the network. The numbers below the layers indicate the number of kernels for convolutional layers, down- or upsampling factor for pooling and upsampling layers, and number of units for fully-connected layers. The displayed numbers are those of the final model used to obtain the presented results for dimensionality reduction to 2 dimensions. For other numbers of dimensions, the only modification made was to change the number of units in the latent representation from 2 to 4, 6, 8 or 10. For the genetic clustering application, the number of units in the latent representation was *k* = 5.

Convolutional layers consist of a number of weight matrices, or kernels, of a specified size. These are used to compute a sliding dot product of the input. Each kernel is convolved over the input sequence along its spatial dimension with a given stride, or step size. In our models we use 1-dimensional convolutional layers with a stride of size 1. The depth of the layer output is thus determined by the number of kernels, and the spatial dimension of the data is unchanged. The pooling layers perform downsampling by applying a max filter over sliding windows of the data, separately for each depth dimension, reducing the size of the spatial dimension and leaving the depth unchanged.

The encoder alternates convolutional and pooling layers to increase the depth and reduce the spatial dimension of the data. The center of the model consists of a series of fully-connected layers in which the latent representation, or encoding, is defined. In contrast to convolutional layers, fully-connected layers contain weights between all pairs of variables in the input and output. The decoder roughly mirrors the structure of the encoder, with upsampling performed by means of nearest-neighbor interpolation to increase the spatial dimension of the data.

Residual connections, shown as black dashed lines in Figure 2, are used to stabilize the training process. These add the output from one layer to later parts of the network, skipping over layers in between, and have been shown to facilitate the optimization of deep networks for different applications (He *et al*. 2015).

Convolutional layers allow the model to capture local patterns and make use of the sequential nature of genetic data, allowing it to incorporate essential features such as LD at various length scales.

In order to facilitate the learning of global patterns in the input data, the model has two additional sets of variables. Each of these sets contains one variable per marker that is updated during the optimization process, allowing the model to capture marker-specific behavior. The two sets of marker-specific variables, illustrated in Figure 2 in red and green, are both inserted into the model by concatenation to layers in the decoder. One set of variables is also concatenated to the model input at every stage of the training process.

The activation function exponential linear unit (elu) is applied to convolutional layers, after which batch normalization is performed. The fully-connected layers also have elu applied to them, except the outermost ones which have linear activation. The final convolution is performed with a kernel size of 1, with linear activation and no batch-normalization. In order to regularize the network and avoid overfitting, dropout is used on the weights of the fully-connected layers, except those surrounding the latent representation, and Gaussian noise is added to the latent representation during training.

Input data is represented as an (*n_samples_* × *n_markers_*) matrix of diploid genotypes, normalized to the range [0,1] by mapping 0 → 0.0,1 →0.5, 2 →1.0. When calculating the loss function, target genotypes are represented using one-hot encoding with 3 classes [*p*(0), *p*(1), *p*(2)] with *p*(*g*) = 1.0 for the true genotype, and *p*(*g*) = 0.0 for the others. Model output *o_ij_* of sample *i* at site *j* has the sigmoid function applied to it, after which it is interpreted as the allele frequency at the site. This scalar is transformed to the format of the targets using the formula for genotype frequencies *f*(*g*) obtained from the principle of Hardy-Weinberg equilibrium:

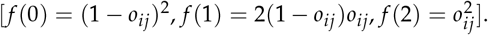

The network is trained to reduce the categorical cross-entropy error (*E*) between target (*y*) and reconstructed (*ŷ*) genotypes, with an added L2 penalty on the values of the latent representation (*e*) for regularization. See Equation 1, where *α* is the regularization factor hyperparameter. Network optimization is performed by means of the ADAM algorithm (Kingma and Ba 2014), with a further exponential decay of the learning rate applied.

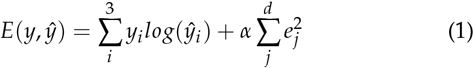

Additional regularization of the training process is performed by means of data augmentation. For every sample in the training process, a fraction of genotypes is randomly set to missing in the input data, represented by the value −1.0. A dimension representing missing and non-missing genotypes, with the values 0 and 1, respectively, is added to the input data, depicted in light blue in Figure 2. The fraction of missingness for each training batch is randomly selected from a pre-defined range. Noise was also added to the input data by introducing incorrect genotypes. With probability 0.2, missing genotypes of a batch were set to random genotype values, drawn from a uniform distribution.

The data was randomly split into training and validation sets consisting of 80% and 20% of samples, stratified by population. The training set was used for network optimization, with the termination criterion that the validation loss had not decreased for 300 epochs.

Different model architecture settings and hyperparameters were evaluated, and the best-performing setups were chosen by means of a hierarchical search procedure. We refer to Supplemental File S1 for details about the evaluated options, including the final model architecture settings and hyperparameter values used for obtaining the presented results.

### Human Origins data set

The data set used to evaluate dimensionality reduction and genetic clustering performance was derived from the fully public present-day individuals of the Affymetrix Human Origins SNP array analyzed in Lazaridis *et al*. (2016). This data set is designed for population genetic studies and represents worldwide genetic variation, containing 2,068 samples from 166 populations. These were categorized into 8 superpopulations, displayed in Figure 1 which also serves as a legend for Figures 3, 5 and 6.

**Figure 3.**
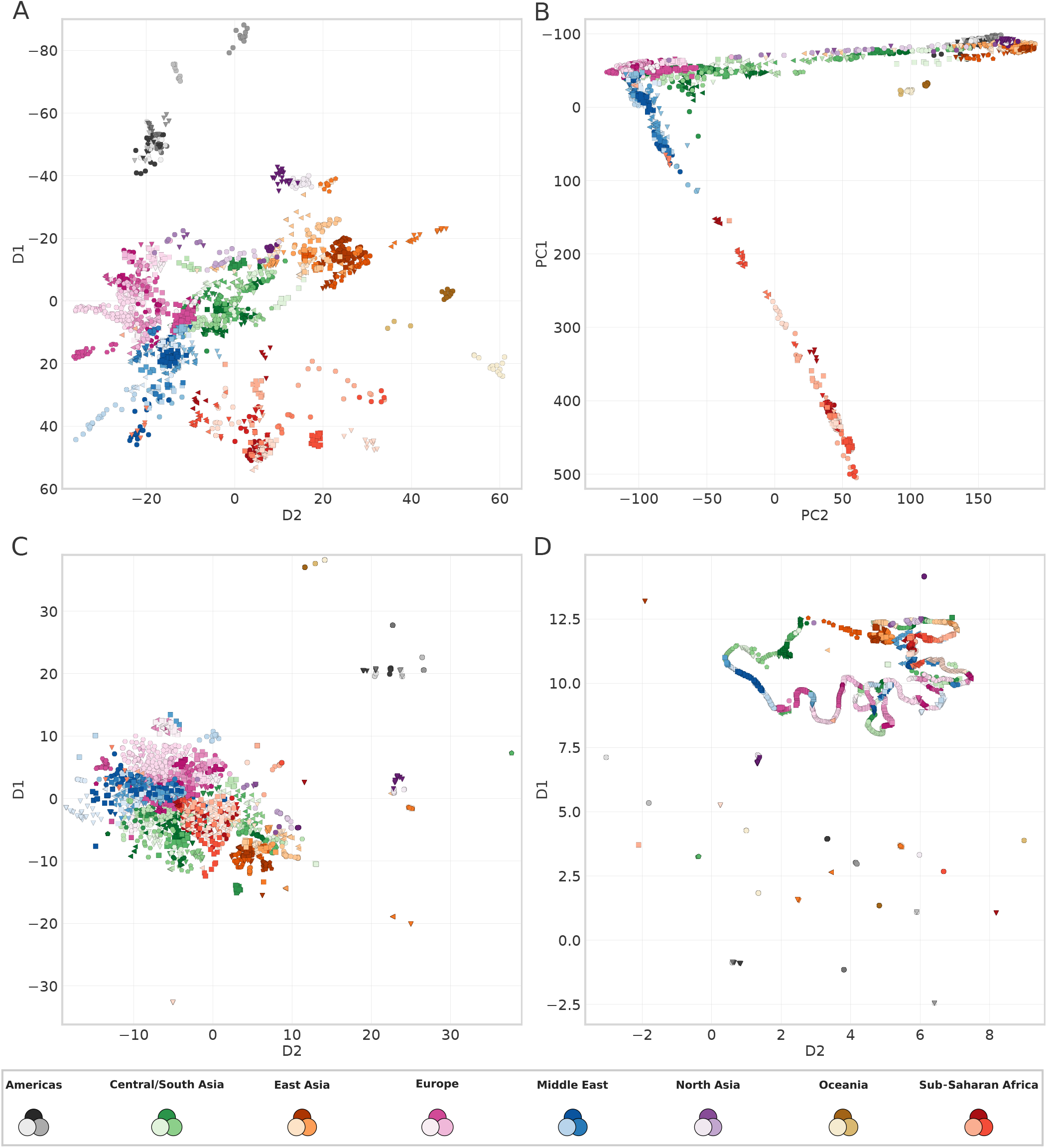
Dimensionality reduction results for GCAE (A), PCA (B), t-SNE (C) and UMAP (D) on the Human Origins data set. For GCAE and PCA, the D1 and PC1 axes have been inverted in order to get a more direct correspondence to the cardinal geographical directions. Legend shows superpopulation colors, full legend with all populations in Figure 1.

**Figure 4.**
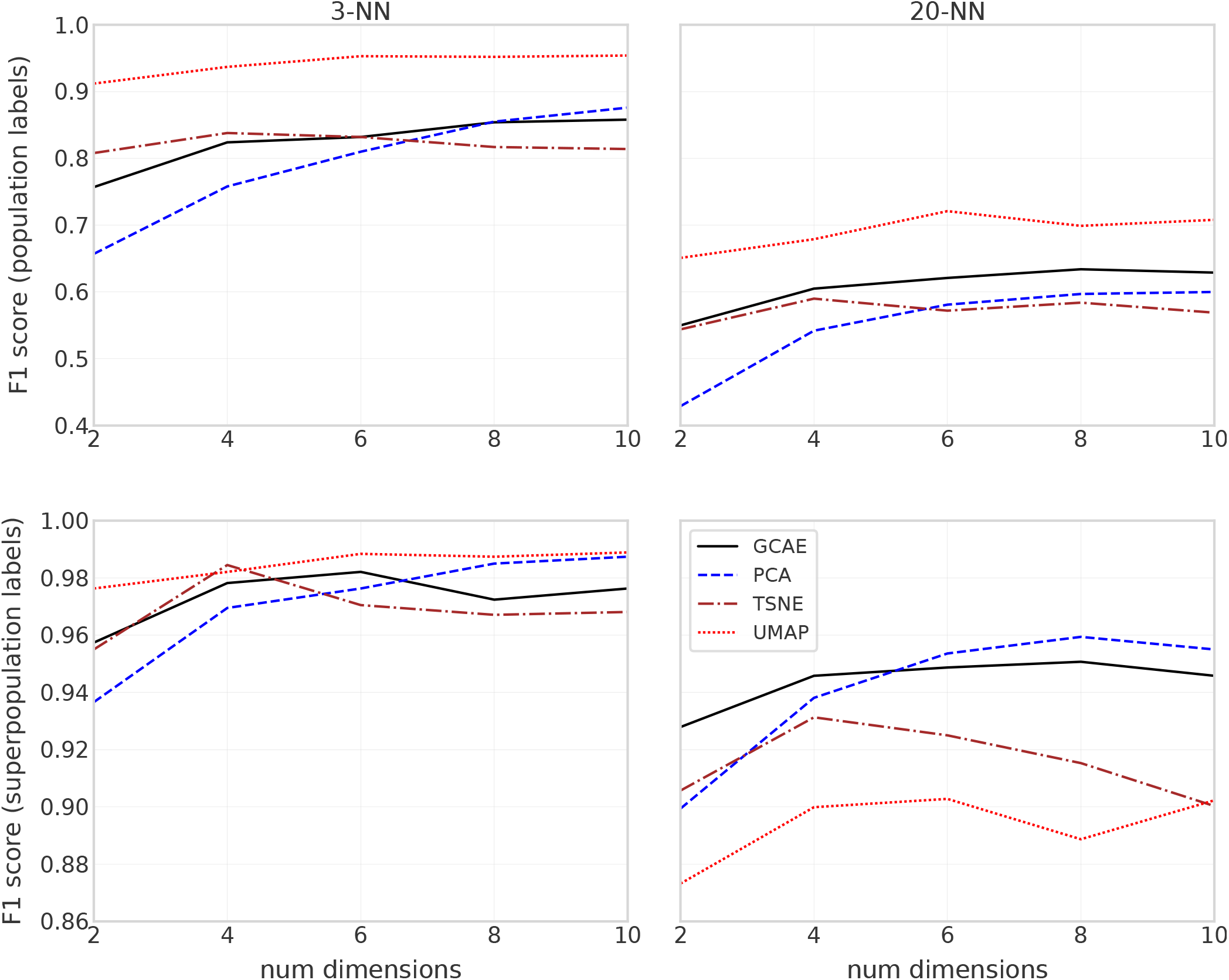
F1 scores for 3-NN and 20-NN classification models based on dimensionality reduction of GCAE, PCA, t-SNE and UMAP using 2-10 dimensions. Top plots show results for using populations as labels in the classification, and bottom ones for using superpopulations as labels.

**Figure 5.**
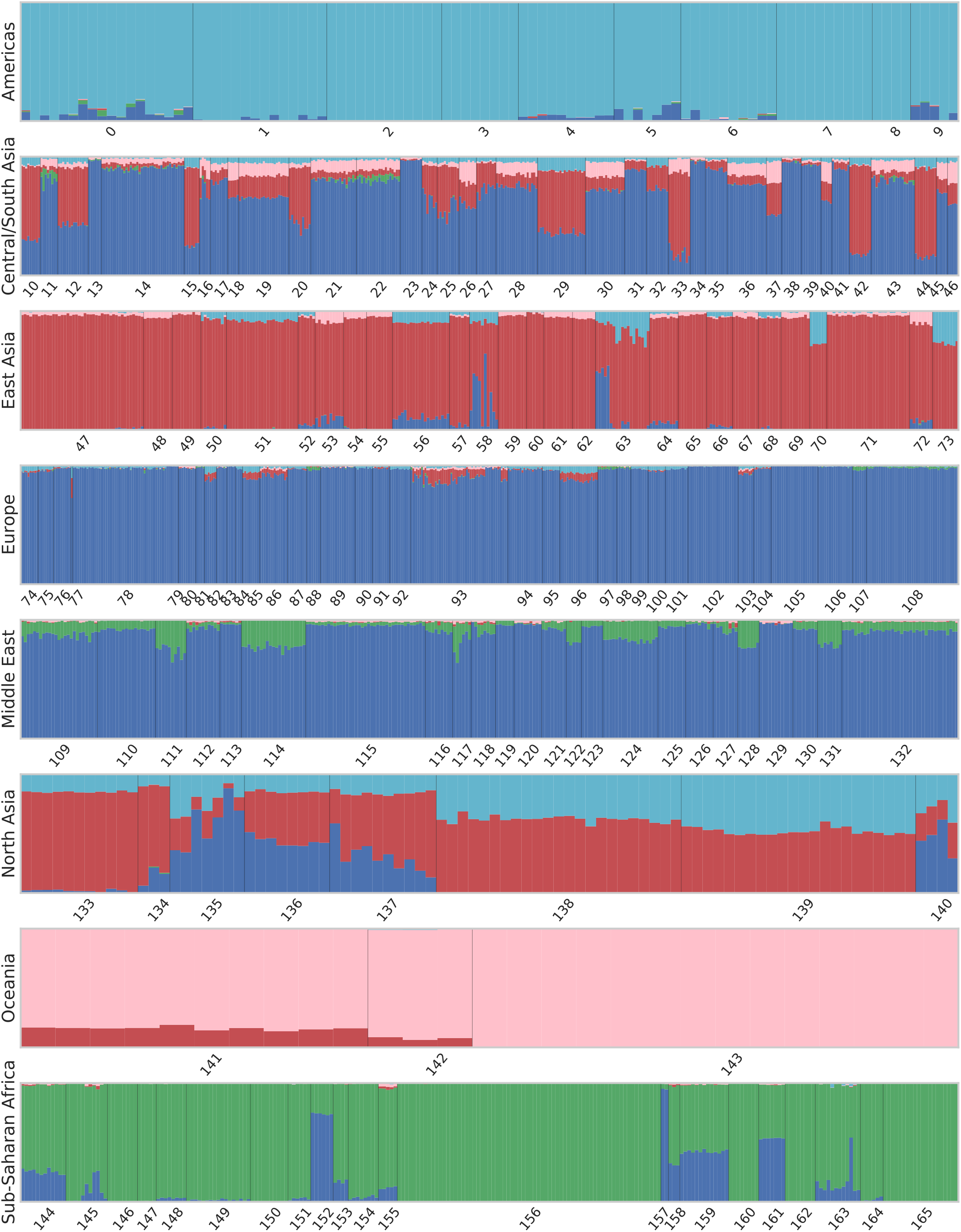
Genetic clustering results with *k* = 5 clusters using ADMIXTURE. Each bar represents a sample from the Human Origins data set, with colors indicating the proportional assignment into *k* clusters for that sample. Samples are ordered by population and superpopulation, with numbering according to the legend in Figure 1.

**Figure 6.**
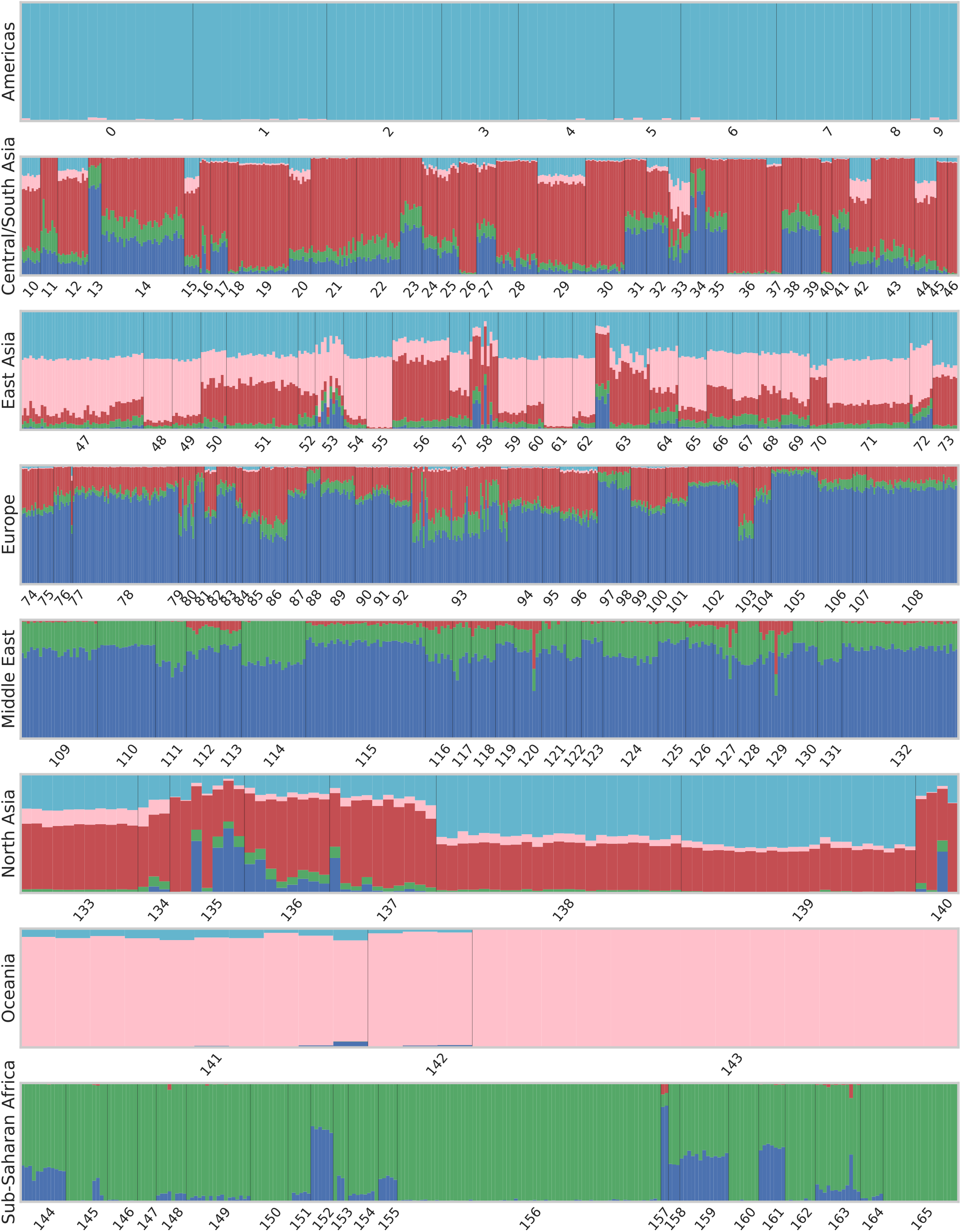
Genetic clustering results with *k* = 5 clusters using GCAE. Each bar represents a sample from the Human Origins data set, with colors indicating the proportional assignment into *k* clusters for that sample. Samples are ordered by population and superpopulation, with numbering according to the legend in Figure 1.

The data was filtered to exclude sex chromosomes and non-informative sites, and one sample (NA13619) was removed due to relation to another (HGDP01382). In order to obtain a single data set for fair comparison between methods, the genotypes were further filtered according to the procedure that is common to perform prior to applying PCA on SNP data. A minor allele frequency (MAF) threshold of 1% was enforced, and LD pruning was performed by removing one of each pair of SNPs in windows of 1 centimorgan that had an allelic R^2^ value greater than 0.2.

As the comparison of robustness of different methods to missing data was beyond the scope of this study, missing genotypes were set to the most frequent value per SNP so as to avoid their influence over dimensionality reduction results. The final data set consisted of 2,067 individuals typed at 160,858 biallelic sites.

### Evaluation of dimensionality reduction performance

Comparison of performance between GCAE, PCA, t-SNE and UMAP was performed by means of evaluating the ability of the dimensionality reduction to capture population structure. A *k*-Nearest Neighbors (*k*-NN) classification model was defined based on the projected data by assigning a population label to each sample based on the most frequent label among its k nearest neighbors.

The evaluation was performed using 2, 4, 6, 8, and 10 latent dimensions. For UMAP, t-SNE and GCAE, hyperparameter tuning was done for each number of latent dimensions, selecting the configuration that yielded the highest F1 score for a 3-NN classification model. See Supplemental File S1 for details. The model with the selected hyperparameters was then used to calculate F1 scores for classification models using 3 and 20 neighbors. The different numbers of neighbors were used to obtain metrics that capture different aspects of performance, e.g. tightness and degree of overlap of the populations clusters.

Further, in order to evaluate the ability of the models to capture global patterns in the data, the models were also evaluated on their ability to classify samples according to membership to the larger continental groups. This was done by calculating the F1 scores using the superpopulations, rather than populations, as labels (see Figure 1).

Classification performance was measured by the F1 score, defined per class label *c* as the harmonic mean of the precision and recall: 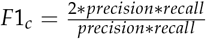. The total F1 score for a model was defined as the average F1 score, weighted by the number of samples per class.

PCA was performed using the same normalization method as in the software SMARTPCA (Patterson *et al*. 2006), i.e. by subtraction of the mean and division with an estimate of the standard deviation of the population allele frequency per SNP. For t-SNE and UMAP, the data was standardized by removing the mean and scaling to unit variance.

The entire data set was used for performance evaluation. This is standard practice for using PCA, t-SNE and UMAP, particularly the latter two as they are non-parametric models. For neural networks, a test set of previously unseen samples is usually used for performance evaluation. In this case, we motivate the use of the entire data set by the nature of the research question itself and the fact that the performance metrics are based on the population labels which are unseen by the network and therefore not used in the model optimization.

### Extension to genetic clustering

The problem of genetic clustering refers to the characterization of individual genomes by proportional assignment to a set of clusters, or genetic components. These may be used in the analysis of population structure and in identifying patterns of genetic variation between populations.

The DL model for genetic clustering was developed by making minor changes to the autoencoder architecture used for dimensionality reduction described above. The number of units in the encoding layer was changed from 2 to *k*, the number of clusters. In order to obtain a proportional assignment, softmax normalization was further applied to the encoding to obtain a vector of *k* values that sum to 1. The loss function was the same as for the dimensionality reduction application, shown in Equation 1. Supplemental File S1 contains more information about the network architecture settings and training options used for the genetic clustering model.

We consider the widely used software ADMIXTURE (Alexander *et al*. 2009) as a comparison method for the genetic clustering application, and present results in a similar manner using bar graphs displaying the proportional assignment of clusters for each sample. For both GCAE and ADMIXTURE, the Human Origins data set described above was used.

### Analysis of output genotypes

In order to further analyze the representation learned by the model, we compared different characteristics of the output genotypes to those of the true ones. First, we studied the distributions of allele frequencies by comparison of the respective site frequency spectra. Secondly, we compared the spatial structure in the two data sets by studying the pattern of LD decay along the chromosome.

For this analysis, we considered a different data set consisting of a single chromosome typed at a denser set of SNPs. As in Battey *et al*. (2020), we used chromosome 22 from the 1000 Genomes phase 3 data (Auton *et al*. 2015), but restricted to biallelic SNPs in the region 24500000-26500000 bp, resulting in 61104 sites.

Network optimization was performed using all 2504 samples, randomly split into training and validation sets consisting of 80% and 20% of samples, stratified by population. The same training procedure as described in was used, as well as a similar 2-D model architecture, with the difference that a larger kernel size was used for the convolutional layers. See Supplemental File S1 for details on the model and hyperparameters evaluated. Model evaluation was done based on the one that yielded the most non-fixed sites for use in the LD calculation. For these experiments, we also used weighting of the loss function to handle the skewed distribution of genotypes, by means of a class-balanced loss based on the effective number of samples (Cui *et al*. 2019) with *β* = 0.95.

LD analysis was performed on the output genotypes of the trained GCAE model, on a subset of samples and SNPs. Similar to the LD analysis performed in Battey *et al*. (2020), we considered samples from the YRI population only, and SNPs in the interval 25000000-26000000 bp. We further applied a MAF threshold of 1% and restricted the genotypes to only include those passing all filters in the ‘strict’ accessibility mask provided by the 1000 Genomes Project.

The R^2^ measure of LD was calculated for the output genotypes and compared to that of the true genotypes. As R^2^ is not defined for pairs of loci where where one or more allele frequencies are equal to zero, sites for which only one allele was present in the output genotypes were excluded from the analysis, in order to consider the same set of sites in the true and output genotypes.

### Implementation

GCAE is implemented in Python 3 using Tensorflow 2 (Abadi *et al*. 2015) and is available at https://github.com/kausmees/GenoCAE as a command-line program. The programs plink 1.9 (Shaun Purcell, Christopher Chang 2020; Chang *et al*. 2015) and bcftools 1.14 (Danecek *et al*. 2021) were used for filtering of genotype data. All other preprocessing of data, as well as performance evaluation and visualization, was implemented in Python. The library scikit-learn 1.0 (Pedregosa *et al*. 2011) was used for calculating the F1 score metric, with scikit-allel 1.3.5 (Miles *et al*. 2021) being used for LD analysis.

PCA and t-SNE were performed using the Python libraries scikit-learn and MulticoreTSNE 0.1 (Ulyanov 2016), respectively. The reference results for genetic clustering were obtained using the software ADMIXTURE 1.3.0 using the em method.

CPU computations were performed on the resources of Uppsala Multidisciplinary Center for Advanced Computational Science (UPPMAX) on a cluster of compute servers equipped with 128 GB memory, each comprising two 8-core Xeon E5-2660 processors. GPU computations were run on National Supercomputer Centre (NSC) at Linköping University, on an NVIDIA SuperPOD with DGX-A100 nodes equipped with 8 NVIDIA A100 Tensor Core GPUs with 40 GB on-board HBM2 VRAM, 2 AMD Epyc 7742 CPUs, 1 TB RAM.

## RESULTS

Figure 3 shows dimensionality reduction results using GCAE, PCA, t-SNE and UMAP on the Human Origins data set. On a global scale, PCA and GCAE result in similar patterns. Both methods are able to capture global geometry to a high degree, with a visible clustering according to superpopulation. Both methods also result in a consistent global pattern with the Sub-Saharan African super-population separating distinctly, a gradient from the Middle East and Europe to South/Central and North Asia onto East Asia that roughly reflects the geography of Eurasia, and a separate Oceanian cluster.

A difference is that while PCA essentially clusters all non-African populations on one dimension, the GCAE plot is more spread out, with a north-south gradient in addition to the east-west relationships mainly captured by PCA. This is visible within superpopulations as well as between them, e.g. by the appearance of the Americas as a distinct cluster and a clearer differentiation of North Asia from and Central/South and East Asia. Populations of the Far East Siberia like Yukagir, Koryak, Chukchi and Eskimo, for example, appear clearly set apart along the D1 axis, which is also the dimension that mainly distinguishes the American samples. Within the African superpopulation, samples from the north and east of the continent are in closer proximity to the Middle Eastern cluster, and differentiated from the cluster of west-African populations. The San populations Khomani and Ju Hoan also separate distinctly, which is not evident in the PCA plot. A similar observation holds for the the Mbuti and Biaka groups of the Congo Basin area.

The neighbor graph-based methods, in contrast, do not display the same global pattern as PCA and GCAE. While t-SNE also shows clustering according to superpopulation, it is a different pattern with Sub-Saharan Africa in the middle, South/Central Asian populations more spread out, and some Asian populations in tight formations at large distances from the main cluster. UMAP shows even more populations appearing in tight and highly separated clusters, with the rest forming a distinct shape on a curved line. While more difficult to interpret, some degree of clustering according to superpopulation is also visible. Overall, both t-SNE and UMAP result in visualizations with less correspondence to global geographical patterns, but a comparatively high degree of clustering of individual populations.

It is important to note that PCA used as a dimensionality reduction method differs from the other methods considered in that there is a choice of which principal components to use. In this part of the evaluation, the focus is on the visual information in the two-dimensional projection, for which it is common to use the first two principal components only. We refrain from discussing other combinations of principal components here, but note that there may be different structure visible when selecting others. For a more complete assessment, we refer to the results of the classification performance below, which indicate the ability of the models to capture population structure when considering multiple dimensions.

Figure 4 shows the F1 scores of 3-NN and 20-NN classification models based on the dimensionality reduction of GCAE, PCA, TSNE and UMAP for 2-10 dimensions. The top and bottom plots show scores for the population and superpopulation classification models, respectively.

For the population classification model, UMAP resulted in the highest F1 scores, which is consistent with the tight clustering of individual populations in Figure 3. t-SNE tends to give relatively high classification performance for lower dimensions, and does not show much improvement in score with increased dimensionality, a trend that is also visible for UMAP. PCA and GCAE show a similar pattern of increasing performance, with GCAE tending to have higher scores.

For the superpopulation classification model, the relative performance of PCA and GCAE to the other models increases. This indicates that the neighbor graph-based methods have less ability to capture global structures, and is particularly evident in the 20-NN model for which they have significantly lower scores. PCA shows quite consistent performance gains for increased dimensionality, indicating that the additional PCs do add structure that is useful for the classification model. For lower numbers of dimensions, GCAE tends to have higher scores than PCA, with performance dropping off for higher dimensions. A possible explanation for this is that regularizing the latent space of GCAE becomes more difficult with increased size, and further exploration of hyperparameter space and regularization methods might be needed to utilize the space to a larger extent.

It is worth noting that our optimization criteria for hyperparameters clearly favors local performance. Both t-SNE and UMAP are models for which hyperparameters control the balance between attention given to local and global aspects of the data, and the authors of UMAP argue that their model can achieve a higher degree of preservation of global patterns in comparison to t-SNE (McInnes *et al*. 2020). The hyperparameter selection process here does not necessarily reflect that of a user interested in using t-SNE or UMAP for visualization purposes, where a more subjective evaluation would most likely be used. To give a more comprehensive view, we evaluate the visualization and clustering results of other hyperparameter combinations for UMAP in Supplemental File S1.

Figures 5 and 6 show the genetic clustering results on the Human Origins data set with 5 clusters using ADMIXTURE and GCAE, with the order of the clusters adjusted to be analogous for comparison. Both models result in distinct American, Oceanian and African clusters, with the latter also showing a very similar pattern of the blue component. The ADMIXTURE results further show the East Asian and European superpopulations as largely distinct clusters, whereas GCAE reveals these as composite.

For European populations, the most prominent components in the GCAE clustering are blue, which is mainly present in the Middle East for both methods, and red. A possible interpretation is that the red cluster signifies the genetic component of herders that migrated to Europe from the Pontic–Caspian steppe around 4.5 kyr ago (Nielsen *et al*. 2017; Haak *et al*. 2015). This would be consistent with a presence in most European populations, with the exception of Sardinians (Lazaridis *et al*. 2014), as well as in South Asia, with particular prominence in e.g. the Kalash (Lamnidis *et al*. 2018; Pathak *et al*. 2018). This component also appears in East Asian populations, which also include American and Oceanian ancestry.

Results for the comparison of output data from GCAE to the corresponding true genotypes are shown in Figure 7. The left plot shows that the GCAE-generated genotypes follow the true distribution of allele frequencies in the population. A tendency to underestimate the presence of the derived allele for low-frequency sites, particularly for values below 0.1 in the true data, is however visible, indicating that rare variation is more difficult for GCAE to capture. Another trend visible in the middle of the frequency spectrum is that output genotypes tend to have a higher presence of the derived allele where the true frequency is below 0.5 and a corresponding underestimation of the derived allele where it is in majority in the true data. This is likely an effect of the balancing performed on the loss function to handle uneven classes, and may be mitigated by finer tuning of the *β* parameter.

**Figure 7.**
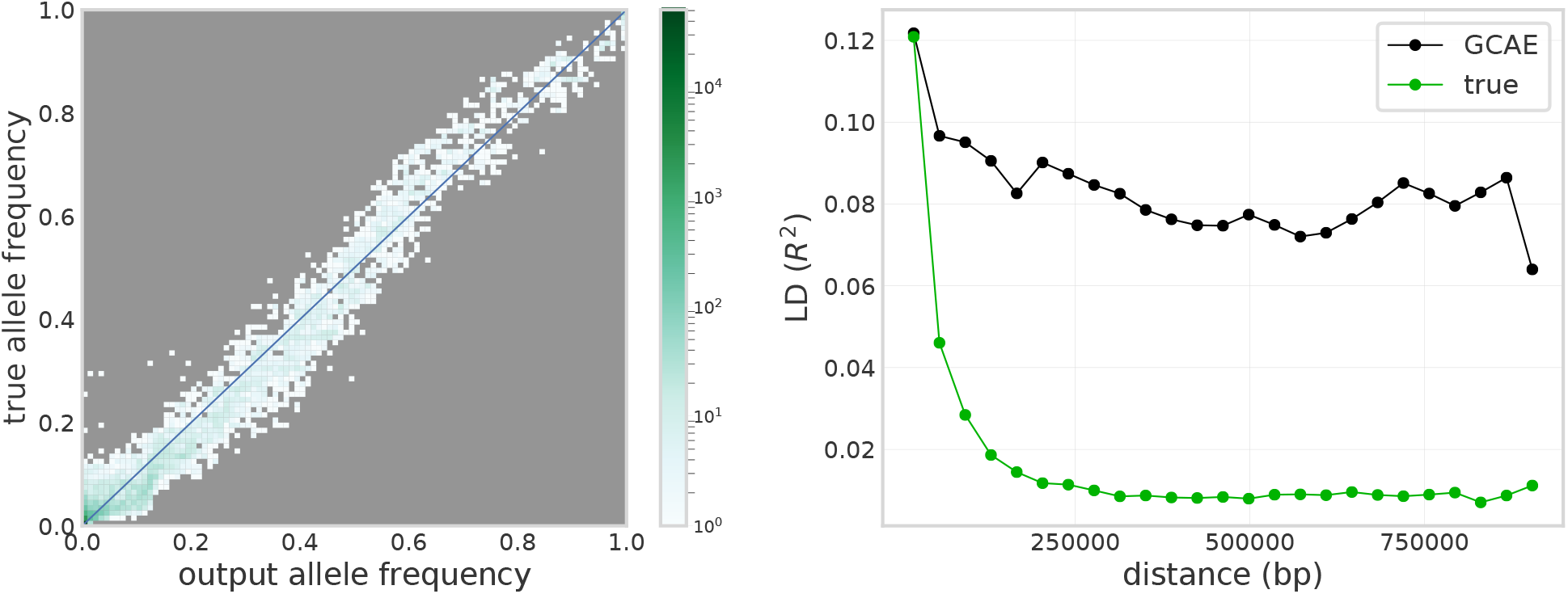
Comparison of output from GCAE to the corresponding true genotypes for the 1000 Genomes chromosome 22 data set. Left: Site frequency spectrum of true genotypes vs. output genotypes, with color indicating number of sites in 100 allele frequency bins. Right: Decay of LD with distance along the chromosome for a subset of samples and SNPs, displaying the mean value of R^2^ for pairs of sites in 25 distance bins.

The right plot shows that output genotypes do show a pattern of decay of LD with distance along the chromosome that reflects that of the true data, with very similar correlation values for the lowest SNP distances. Although the correlation between sites tends to be higher in the GCAE-generated data for longer distances, with a less smooth decay curve, the results suggest that GCAE is able to define the coding into the latent space in a way that takes local spatial structure into account.

## DISCUSSION

One approach to evaluation of dimensionality reduction involves assessing the correspondence between transformed data and its geographical sampling location. Quantitative studies have shown that geographical effects in the form of migration and the impact of physical distance on gene flow play a role in creating population structure (Wang *et al*. 2012). For PCA, striking similarities to geography have mainly been reported from limited geographical areas such as Europe (Novembre *et al*. 2008; Lao *et al*. 2008), with world-wide cohorts generally resulting in a less resolved V-shape similar to that shown in Figure 3 B (Jakobsson *et al*. 2008; Biswas *et al*. 2009). As previously mentioned, t-SNE and UMAP tend to focus on local relationships, although Diaz-Papkovich *et al*. (2021) discusses that for UMAP, careful filtering of the data can cause geographical features to be highlighted more, but again, mainly when applied to relatively homogeneous data sets. The Human Origins data set considered here represents worldwide genetic variation, and the visualization results show that GCAE displays robustness to this high degree of diversity, yielding a representation that reflects global geographical patterns.

The interpretation of dimensionality reduction results, particularly the inference of the underlying processes behind observed structure, is however not always straightforward. The clustering of samples in reduced-dimensional space can reflect characteristics at various scales in the data, ranging from the presence of a particular variant to continental ancestry. In Novembre and Stephens (2008) and François *et al*. (2010), for example, the effects of past migration and expansion events on PCA is discussed, and how the assumptions of linearity and orthogonality of the model can result in counter-intuitive patterns in PC-space. The effects of attributes of the data such as LD and so-called “informative missingness”, due to e.g. different sequencing panels or the higher uncertainty associated with heterozygote calls, on dimensionality reduction are also extensively discussed in Patterson *et al*. (2006).

The F1 score of a classification model based on the dimensionality reduction is not a simple metric for which the method with the highest score is the most correct. Nonetheless, the results indicate relevant, systematic differences between the models. With respect to the performance metric considered, PCA and GCAE both seem to be able to make more efficient use of increasing numbers of dimensions in the latent space than t-SNE and UMAP. Further, the three nonlinear methods tend to reveal more fine-scale patterns and yield a more resolved representation than PCA, at least for lower number of dimensions. A key difference, however, is that GCAE takes a more global approach that preserves the meaning of distances between clusters to a larger extent than the neighbor graph-based methods.

Other deep-learning based methods that generate genotypes have shown varying capabilities to preserve spatial properties in terms of decay of LD with distance along the chromosome. The variational autoencoder of Battey *et al*. (2020) failed to reproduce LD decay, while both the generative adversarial network (GAN) and RBM models of Yelmen *et al*. (2021) captured LD patterns well, with correlation at larger distances in particular preserved to a higher extent than for GCAE. An important methodological difference between GCAE and these other models, and also PCA, t-SNE and UMAP, is that they do not take the sequential nature of genotype data into account. Rather, they treat every SNP as an independent variable, whereas convolution treats relationships between nearby sites differently than any arbitrary pair of variants. We show that convolutional architectures can be used to capture local spatial patterns, and believe that additional convolutional and max-pooling layers can improve LD accuracy over longer distances.

Our results demonstrate that the use of convolution is feasible for genotype data in spite of its fundamental differences to images, which such networks are typically applied to. Genotype data is position-dependent, with a unique meaning to every dimension. Pixels in images, in contrast, typically represent information that is translation invariant. Our experiments indicated that the incorporation of positional information in the form of marker-specific variables that the model can optimize during the training process improved performance. We therefore suggest this as a means of allowing the model to represent some of the global information that is lost with convolution. An additional, more explicit, method to include sequential information would be to include e.g. genetic distance or position as part of the input data, as done in Chan *et al*. (2018);Adrion *et al*. (2020), which we leave for future work.

We also note that in order to obtain a fair comparison, we have performed filtering of the SNP set in terms of MAF and LD according to standard protocols for PCA even though these steps are not necessarily required for GCAE, in which such patterns can be learned during the training process. The GCAE architecture also includes a representation of missing genotypes, unlike the other models. In practice, missing data is often handled by either imputation with the empirical mean and/or filtering to remove sites with high missingness. In this sense, GCAE can present a more robust alternative that is more suitable for low-coverage samples such as ancient DNA, requiring less filtering and allowing more of the data to be retained for analysis.

Many commonly used methods for genetic clustering such as TreeMix (Pickrell and Pritchard 2012), ADMIXTUREGRAPH (Lep-pälä *et al*. 2017) and GLOBETROTTER (Hellenthal *et al*. 2014) are, unlike the dimensionality reduction methods discussed, model-based. STRUCTURE (Pritchard *et al*. 2000), for example, represents LD and includes explicit modeling of admixture blocks and the transitions between them. The model assumes the existence of a set of differentiated ancestral populations, and that the sample is a result of relatively recent mixing of these. ADMIXTURE, which we use as a reference method in this work, is based on the same underlying statistical model.

GCAE, in contrast, constitutes a more flexible and data-driven approach, which may be an advantage in scenarios where the data does not conform to explicit modeling assumptions. Our results demonstrate that GCAE is able to capture very similar population structure as that found by ADMIXTURE, while also identifying additional characteristics for some populations that are consistent with existing findings in the literature.

As previously discussed regarding dimensionality reduction, interpretation and evaluation of genetic clustering is not straight-forward. The correctness of an assignment is not well-defined, and different underlying processes can give rise to similar observed patterns (Lawson *et al*. 2018). Evaluation of results requires additional information, such as putting them into the context of methods based on different underlying models, and the analysis of metrics like F and D statistics.

A purely data-driven black-box approach such as DL can be difficult to interpret. The features used in the data transformation are unknown and therefore cannot be used for validation of whether certain modeling assumptions hold for the data in question. On the other hand, the alternative methodology of GCAE allows it to capture additional aspects of the data, and therefore provide a useful complement to the toolset used for exploratory data analysis in population genetics.

Another characteristic of convolutional layers is that they require less trainable variables than a corresponding fully-connected layer, leading to reduced computational requirements for training. Depending on overall network architecture and training strategy, this may allow for the design of models that are more feasible to train on large data sets.

When running on CPUs, using 11 cores on UPPMAX, training of the dimensionality reduction models took between 26.9 and 99.8 hours. Using the GPU on NSC, times ranged between 1.5 and 5.9 hours. The genetic clustering model took 46.6 hours on CPU, and the model trained on 1000 Genomes data for which output genotypes were analyzed took 45 minutes on GPU. As a comparison, PCA took 11 minutes, and t-SNE and UMAP ranged between 20 minutes and 4 hours on the CPU setup on UPPMAX.

The computational requirements of GCAE are thus greater than that of the other models, although the use of GPUs can improve performance significantly. As the purpose of this study is to evaluate the applicability of convolutional autoencoders to the chosen problems, optimization of computational efficiency is considered out of scope and left for future work.

Our results demonstrate that GCAE can learn features that characterize genotype data in a meaningful way. The minor model changes required to change the application from dimensionality reduction to genetic clustering further demonstrate the flexibility of the method, and future efforts will involve investigating the application of GCAE to other problems. A simple alternative application would be imputation of missing genotypes. As the training procedure is based on reconstructing the input, and since we already include a representation of missing data, this would mainly involve finding a suitable number of units to use in the latent layer. The model can also be used for generation of artificial genotypes by entering data into the decoder that does not correspond to the encoding of an empirical sample. This can be done e.g. by perturbing the encoding of actual individuals or selecting values from a specific part of latent space. If the space is regular enough, one could use the clustering it has defined to simulate samples from a particular population or some other characteristic property learned by the model, e.g. ancient data. We are also currently exploring the use of GCAE in the context of quantitative genetics by incorporating phenotypic information into the model.

## DATA AVAILABILITY

The fully public Affymetrix Human Origins present-day individuals from Lazaridis *et al*. (2016) are available for download from https://reich.hms.harvard.edu/datasets. The 1000 Genomes phase 3 data set is available at https://www.internationalgenome.org/data-portal/data-collection/phase-3.

## ACKNOWLEDGMENTS

The authors acknowledge the use of computational resources provided by Swedish National Infrastructure for Computing (SNIC) and associated centers under projects Berzelius-2021-30, UPPMAX 2020/2-5, SNIC 2019/8-38 and SNIC 2020/5-91. CN also acknowledges funding by Formas (grant numbers 2017-00453, 2020-00712).

