## Supplemental File S1 for "A deep learning framework for characterization of genotype data"

#### Optimization of model design and training of GCAE

The fundamental structure of the model was chosen based on empirical experience from previous experiments (not published) and related literature. This includes the types of layers used (convolutional, maxpooling, fully-connected, and upsampling by means of nearest-neighbor interpolation) and the ordering of these. We had also found benefits of the use of residual connections and dropout for the weights of fully-connected layers, except those on either side of the latent representation. Previous experience had also shown that the use of the two sets of marker-specific variables had improved model performance, and indicated suitable locations to insert them in the decoder.

Based on this fundamental structure, an evaluation of different architecture settings, shown in Table 1, was performed. These tests were done using a limited set of hyperparameter combinations. Once the final model architecture and settings had been decided, a larger search among the hyperparameter values shown in Table 2 was performed. We note that not all combinations of settings in the two tables were evaluated for the different applications.

Several additional strategies for improving the training process were also evaluated throughout the above described process. We found that using a linear activation function on the outermost fully-connected layers led to a more stable training that was less likely to collapse and avoided artefacts in the latent representation. Data-augmentation by means of setting input genotypes to missing was found to reduce overfitting. This was implemented by randomly setting $f$ genotypes to missing in each batch, where $f$ was randomly selected from $[0.0, 0.1, 0.2, 0.3, 0.4]$. Introducing errors with a probability of 0.2 by setting all missing values in a batch to genotype values drawn from a uniform distribution was also found to improve performance.

| Parameter | Values |
|---|---|
| kernel size convolution | 3, 5, 10 |
| pool size maxpool | 2, 3, 5 |
| stride maxpool | 2, 3 |
| number of filters/kernels convolution | 8, 16, 32 |
| dropout rate | 0.0, 0.01, 0.1 |
| number of units in fully-connected layers | 15, 25, 50, 75 |

Table 1: Model architecture settings

| Parameter | Values |
|---|---|
| learning rate | 3.2e-06, 1e-05, 3.2e-05, 0.0001, 0.00032, 0.001, 0.0032, 0.01 |
| exponential decay rate | 0.92, 0.94, 0.96, 0.98 |
| decay every epochs | 10, 20 |
| regularization factor | 3.2e-09, 1e-08, 3.2e-08, 1e-07. 3.2e-07, 1e-06, 3.2e-06 |
| noise std | 10.0, 3.2, 1.0, 0.32, 0.1, 0.032, 0.01 |
| batch size | 30, 50, 60, 75, 100 |

Table 2: Hyperparameters evaluated for GCAE

### Final implementations

Table 3 shows the architecture settings and hyperparameters used to obtain the presented dimensionality reduction (DR) and genetic clustering (GC) results, as well as the model selected for analysis of output genotypes (OA). For the DR results with more than 2 dimensions, the best model was selected based on evaluating the F1 score of a 3-NN population classification model. For GC and 2-dimensional DR, selection was done based on the F1 score, genotype concordance of the reconstructed data as well as inspection of the visualization. For OA, the model that yielded the highest number of non-fixed sites for LD analysis was selected. In all cases, the epoch at which to stop training was selected as the one where the validation loss was minimized, using a patience of 300 epochs.

| Parameter | DR-2 | DR-4 | DR-6 | DR-8 | DR-10 | GC | OA |
|---|---|---|---|---|---|---|---|
| kernel size convolution | 5 | 5 | 5 | 5 | 5 | 5 | 10 |
| number of filters/kernels convolution | 8 | 8 | 8 | 8 | 8 | 8 | 8 |
| pool size maxpool | 5 | 5 | 5 | 5 | 5 | 5 | 5 |
| stride maxpool | 2 | 2 | 2 | 2 | 2 | 2 | 2 |
| dropout rate | 0.01 | 0.01 | 0.01 | 0.01 | 0.01 | 0.01 | 0.01 |
| number of units in fully-connected layers | 75 | 75 | 75 | 75 | 75 | 75 | 75 |
| learning rate | 0.01 | 0.00032 | 0.00032 | 1.00e-05 | 1.00e-05 | 0.001 | 0.0001 |
| exponential decay rate | 0.92 | 0.92 | 0.92 | 0.98 | 0.98 | 0.94 | 0.98 |
| decay every epochs | 10 | 10 | 10 | 20 | 20 | 10 | 20 |
| regularization factor | 1e-08 | 1e-06 | 1e-06 | 1e-07 | 3.2e-08 | 1e-08 | 3.2e-09 |
| noise std | 0.032 | 0.32 | 1.0 | 3.2 | 10 | 0.032 | 3.2 |
| batch size | 60 | 60 | 60 | 30 | 30 | 60 | 30 |

Table 3: Final architecture settings and hyperparameters used for the dimensionality reduction (DR), genetic clustering (GC) and output analysis (OA) models.

### Hyperparameter tuning of t-SNE

A grid search of learning rate and perplexity was performed for t-SNE, values are shown in Table 2. Default values were used for all other parameters and settings. The values selected, for which results are shown in the main text, are shown for each number of dimensions

| parameter | values |
|---|---|
| learning rate | 43, 170, 200, 225, 250, 300, 350, 400, 450, 500, 650, 600 |
| perplexity | 5, 10, 20, 30, 45, 50, 75,100, 125, 175, 200 |

Table 4: Hyperparameters evaluated for t-SNE.

### Hyperparameter tuning of UMAP

A grid search of the hyperparameters number of neighbors, minimum distance, and spread was performed for UMAP for 2,4,6,8 and 10 number of dimensions. The values considered are shown in Table 6. Default values were used for all other parameters and settings. Optimal hyperparameters were selected based on the most accurate resulting 3-NN population classification model, measured using the F1-score. Table 7 shows the selected hyperparameters for UMAP for the different numbers of dimensions.

| number of dimensions | learning rate | perplexity |
|---|---|---|
| 2 | 300 | 10 |
| 4 | 450 | 175 |
| 6 | 43 | 20 |
| 8 | 650 | 45 |
| 10 | 43 | 50 |

Table 5: Optimal hyperparameters for t-SNE for different number of dimensions, selected based on highest F1 score for a 3-NN population classification model.

| parameter | values |
|---|---|
| number of neighbors | 3, 5, 10, 20, 50, 100, 200 |
| minimum distance | 0.05, 0.1, 0.25, 0.5, 0.75, 1.0 |
| spread * | 0.05, 0.1, 0.25, 0.5, 0.75, 1.0 |

Table 6: Hyperparameters evaluated for UMAP.
*The implementation used requires spread to be larger than or equal to minimum distance, so only such combinations were considered.

In [1], number of neighbors is described as influencing the tradeoff between fine-grained and large-scale manifold features, with smaller values resulting in detailed structure being captured at a loss of "big picture" information, and larger values keeping more large-scale manifold structures. Minimum distance is explained as affecting how closely points end up to each other in the output, where low values result in densely packed points, with manifold structure more likely to be accurately represented. In the code documentation of the implementation used, spread is described as the effective scale of the points, which in combination with minimum distance determines the degree of aggregation of points into clusters.

Thus, there is a tradeoff between local and global behavior for UMAP that can be controlled using the hyperparameters. Our optimization criteria favor local behavior, which is visible in the results in the main text. UMAP outperformed the other methods for the population classification model, but not for the superpopulation classification model, with particularly poor comparative performance for the 20-NN model.

In order to get a more comprehensive view of the behavior of UMAP, we also evaluate its performance for different hyperparameter values. For this, we consider reduction to 2 dimensions, and select hyperparameter values that cover the ranges considered for number of neighbors, minimum distance and spread. Table 8 shows the model identifiers and hyperparameter combinations considered. Also shown are two metrics of classification performance: F1 scores for a 3-NN population classification model and a 20-NN superpopulation classification model, reflecting local and global clustering performance. Also shown in the table are the corresponding metrics for the GCAE, PCA and t-SNE models for

| dimensions | num neighbors | min distance | spread |
|---|---|---|---|
| 2 | 3 | 0.05 | 0.25 |
| 4 | 3 | 0.05 | 0.25 |
| 6 | 3 | 0.1 | 0.1 |
| 8 | 3 | 0.05 | 0.05 |
| 10 | 3 | 0.1 | 0.1 |

Table 7: Optimal hyperparameters for UMAP for different number of dimensions, selected based on highest F1 score for a 3-NN population classification model.

2 dimensions from the main article text. Figures 1 - 3 show the dimensionality reduction results for the UMAP models.

It is visible that the hyperparameters have a large effect on the resulting dimensionality reduction. Increasing the number of neighbors decreases the tendency towards small, isolated clusters, and lower minimum distance and spread give more densely packed points. The F1 score of the 3-NN population classification model tends to be higher for lower neighbor counts, which is expected as those values promote local structures being captured. The more global performance, as measured by F1 score for a 20-NN superpopulation classification model, has less of a clear trend and seems to be less affected by the hyperparameter values. All models considered in Table 8 had lower F1 scores for this metric than GCAE, t-SNE and PCA. The results thus confirm the tradeoff between local and global performance for UMAP, whereas we found that for GCAE, classification of superpopulations tended to improve with that of populations, suggesting at a qualitative difference between the two methods.

We also note that the UMAP projections differ from those of PCA and GCAE in terms of the relative positions of the superpopulations. For the latter two, there is a tendency towards a gradient with Africa and East Asia at opposite ends, with Middle Eastern, European and South/Central Asian populations between them, roughly in this order. This pattern is evident in the PCA plot, and we also noted that GCAE results consistently showed the same tendency when performing the hyperparameter optimization. In the UMAP plots, the African and East Asian clusters tend to appear closer together, often with the European cluster at the extreme end of the larger collection of clusters. This suggests systematic differences in the nature of the global patterns captured by the dimensionality reduction between UMAP and the other methods.

### References

[1] Leland McInnes, John Healy, and James Melville. *UMAP: Uniform Manifold Approximation and Projection for Dimension Reduction.* 2020. arXiv: `1802.03426 [stat.ML]`.

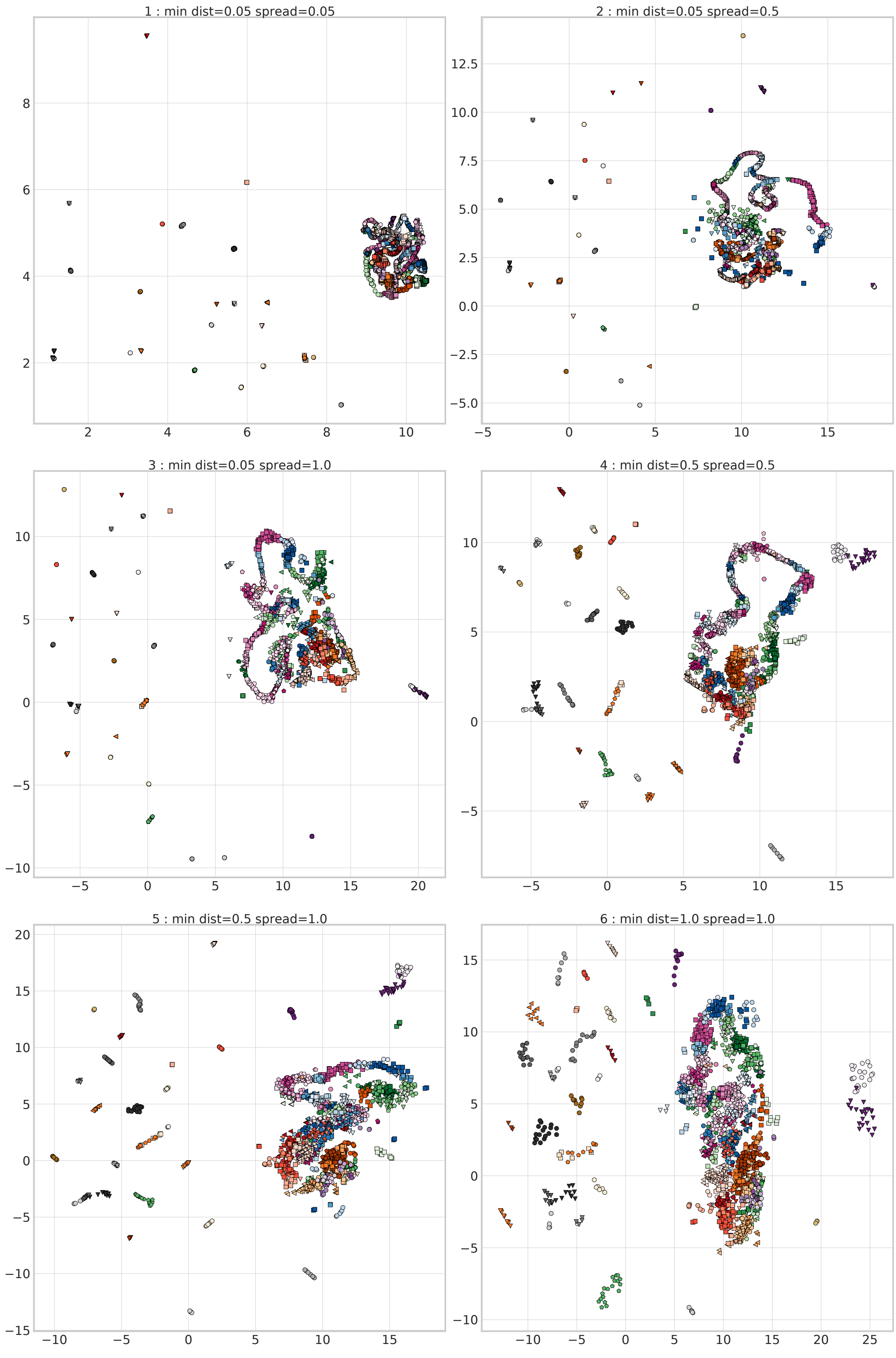


Figure 1: Dimensionality reduction results of UMAP models with number of neighbors = 3.

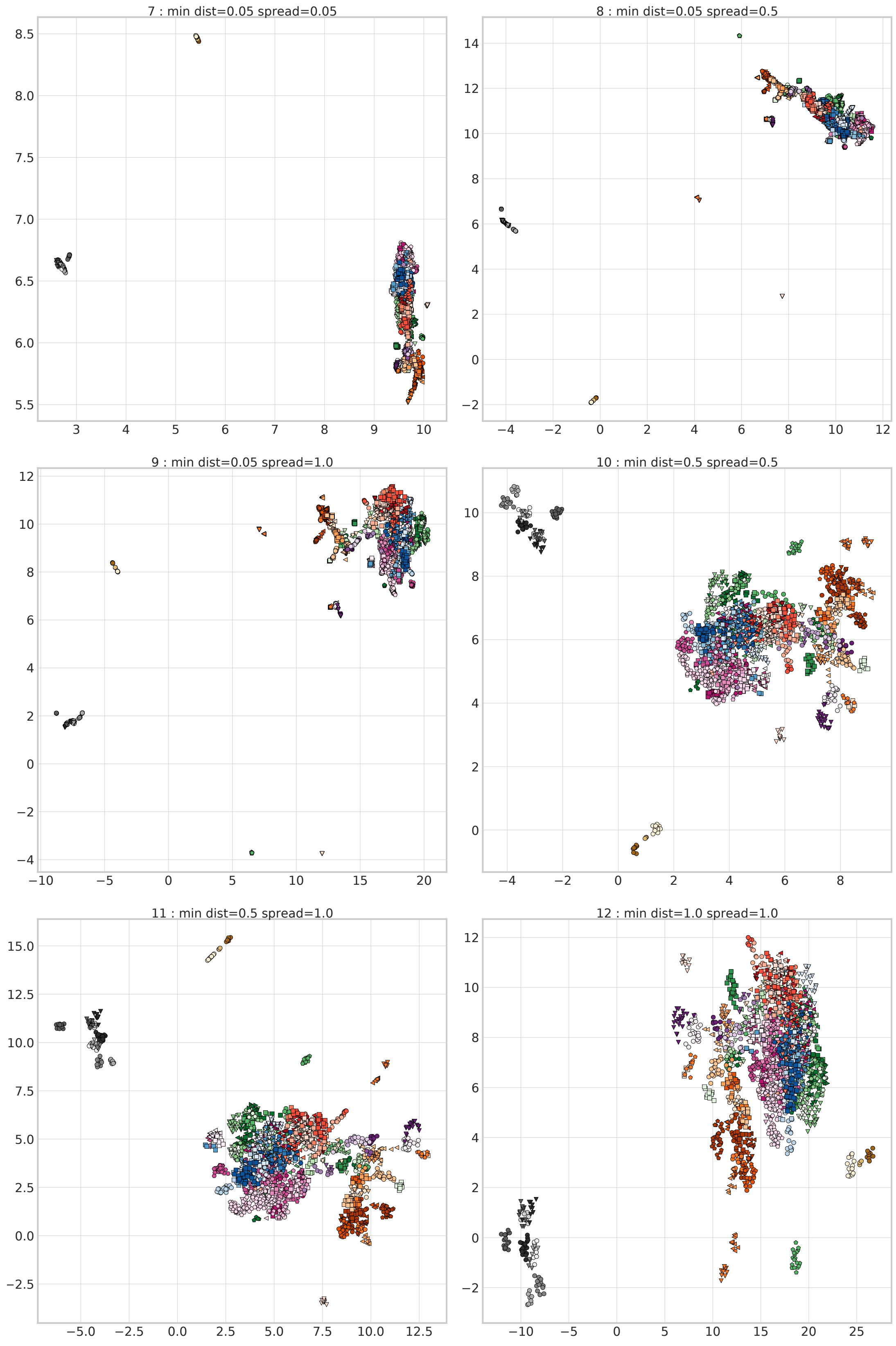


Figure 2: Dimensionality reduction results of UMAP models with number of neighbors = 20.

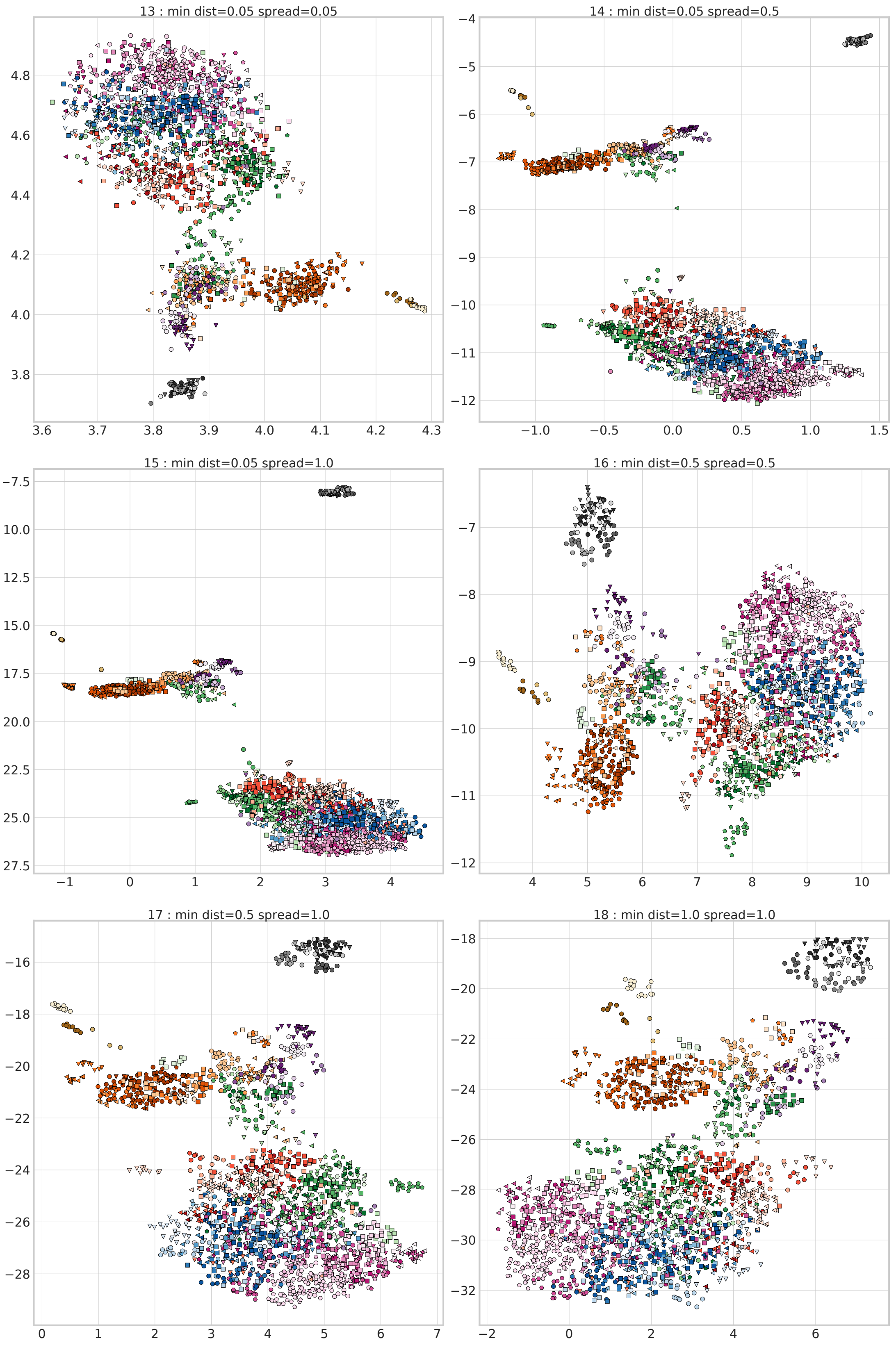


Figure 3: Dimensionality reduction results of UMAP models with number of neighbors = 200.

| model id | num neighbors | min distance | spread | F1 score 3-NN (population labels) | F1 score 20-NN (superpopulation labels) |
|---|---|---|---|---|---|
| 1 | 3 | 0.05 | 0.05 | 0.847 | 0.825 |
| 2 | 3 | 0.05 | 0.5 | 0.853 | 0.798 |
| 3 | 3 | 0.05 | 1.0 | 0.786 | 0.828 |
| 4 | 3 | 0.5 | 0.5 | 0.814 | 0.857 |
| 5 | 3 | 0.5 | 1.0 | 0.719 | 0.801 |
| 6 | 3 | 1.0 | 1.0 | 0.71 | 0.827 |
| 7 | 20 | 0.05 | 0.05 | 0.6 | 0.777 |
| 8 | 20 | 0.05 | 0.5 | 0.726 | 0.857 |
| 9 | 20 | 0.05 | 1.0 | 0.709 | 0.861 |
| 10 | 20 | 0.5 | 0.5 | 0.659 | 0.833 |
| 11 | 20 | 0.5 | 1.0 | 0.658 | 0.849 |
| 12 | 20 | 1.0 | 1.0 | 0.636 | 0.828 |
| 13 | 200 | 0.05 | 0.05 | 0.401 | 0.734 |
| 14 | 200 | 0.05 | 0.5 | 0.555 | 0.815 |
| 15 | 200 | 0.05 | 1.0 | 0.567 | 0.836 |
| 16 | 200 | 0.5 | 0.5 | 0.529 | 0.814 |
| 17 | 200 | 0.5 | 1.0 | 0.553 | 0.829 |
| 18 | 200 | 1.0 | 1.0 | 0.529 | 0.814 |
| GCAE | - | - | - | 0.757 | 0.928 |
| t-SNE | - | - | - | 0.808 | 0.906 |
| PCA | - | - | - | 0.657 | 0.899 |

Table 8: UMAP models with different hyperparameter combinations, and their F1 scores for two classification models based on reduction to 2 dimensions. The last three rows show the same F1 score metrics for the GCAE, PCA and t-SNE models for 2 dimensions from the main article text.
